# M^2^IA: a Web Server for Microbiome and Metabolome Integrative Analysis

**DOI:** 10.1101/678813

**Authors:** Yan Ni, Gang Yu, Huan Chen, Yongqiong Deng, Philippa M Wells, Claire J Steves, Feng Ju, Junfen Fu

## Abstract

**Background:** In the last decade, integrative studies of microbiome and metabolome have experienced exponential growth in understanding their impact on human health and diseases. However, analyzing the resulting multi-omics data and their correlations remains a significant challenge in current studies due to the lack of a comprehensive computational tool to facilitate data integration and interpretation. In this study, we have developed a microbiome and metabolome integrative analysis pipeline (M^2^IA) to meet the urgent needs for tools that effectively integrate microbiome and metabolome data to derive biological insights.

**Results:** M^2^IA streamlines the integrative data analysis between metabolome and microbiome, including both univariate analysis and multivariate modeling methods. KEGG-based functional analysis of microbiome and metabolome are provided to explore their biological interactions within specific metabolic pathways. The functionality of M^2^IA was demonstrated using TwinsUK cohort data with 1116 fecal metabolites and 16s rRNA microbiome from 786 individuals. As a result, we identified two important pathways, i.e., the benzoate degradation and phosphotransferase system, were closely associated with obesity from both metabolome and microbiome.

**Conclusions:** M^2^IA is a public available web server (http://m2ia.met-bioinformatics.cn) with the capability to investigate the complex interplay between microbiome and metabolome, and help users to develop mechanistic hypothesis for nutritional and personalized therapies of diseases.

## Background

The study of microbiome in human health has experienced exponential growth over the last decade with the advent of new sequencing technologies for interrogating complex microbial communities^[1]^. Meanwhile, metabolomics has been an important tool for understanding microbial community functions and their links to health and diseases through the quantitation of dozens to hundreds of small molecules^[2]^. The gut microbiota is considered a metabolic ‘organ’ to protect the host against pathogenic microbes, modulate immunity, and regulate metabolic processes, including short chain fatty acid production and bile acid biotransformation^[3]^. Conversely, these metabolites can modulate gut microbial compositions and functions both directly and indirectly^[4]^. Thus, the microbiota-metabolites interactions are important to maintain the host health and wellbeing. Integrative data analysis of gut microbiome and metabolome can offer deep insights on the impact of lifestyle and dietary factors on chronic and acute diseases (e.g., autoimmune diseases, inflammatory bowel disease^[5]^, cancers^[6]^, type 2 diabetes and obesity^[7]^, cardiovascular disorders, and non-alcoholic fatty liver disease^[8]^), and provide potential diagnostic and therapeutic targets^[9]^.

In the last decade, metabolomics studies in microbiota-related research have increased in a wide range of research areas, such as gastroenterology, biochemistry, endocrinology, microbiology, genetics, to nutrition, food science and pharmacology (Figure S1). However, both metagenomics/16s rRNA-based high-throughput sequencing technologies and mass spectrometry/nuclear magnetic resonance-based metabolomics platforms can produce large and high-dimensional data, posing a major challenge for subsequent data integration^[10, 11]^. Current integration analysis methods mainly focus on statistical correlations between microbiome and metabolome, such as the spearman correlations and partial least squares discriminant analysis (Table S1). Pedersen et al. recently summarized a step-by-step computational protocol from their previous study that applied WGCNA, a dimension reduction method, to measure the correlations among the host phenotype, gut metagenome, and fasting serum metabolome. However, separate analyses of -omics data through one or two statistical methods only provide fragmented information and do not capture the holistic view of disease mechanisms. Moreover,although much bioinformatics work has been done to process and analyze the individual omics data, to date, it still lacks a comprehensive strategy or a computational tool to analyze the correlations between microbiome and metabolome^[12]^. To rapidly advance microbiome and metabolome data integration and understand their roles in diverse diseases, advanced computational methods for multi -omics data integration and interpretation need to be developed^[2]^.

In this study, we have developed a comprehensive computational tool, a standardized workflow for microbiome and metabolome integrative analysis (M^2^IA). M^2^IA combines a variety of univariate and multivariate methods for correlation analysis and data integration. In addition, functional network analysis implemented with KEGG metabolic pathway database can provide deep insights of their biological correlations, i.e., the possible participation of identified microbes in a specific metabolic reaction or metabolic pathway. M^2^IA accepts the microbiome data from 16S rRNA gene or shotgun metagenomic sequencing technologies, and metabolomics data from mass spectrometry or NMR spectroscopy platforms. Other than current bioinformatics tools, M^2^IA is an automated data analysis and reporting pipeline that can be applied by users with little training in bioinformatics. M^2^IA is a web-based server with user-friendly interface, and public available to academic users through http://m2ia.met-bioinformatics.cn.

### Implementation

M^2^IA is developed in Java web system, with html, Javascript technology implemented for interactive interface and JFinal and Mysql databases for data management in the backend. All the statistical analyses and visualization were implemented in R programming language. The entire system is deployed on a cloud server with 16 GB of RAM and four virtual CPUs with 2.6 GHz each. Users can register an academic account to manage their own research projects. M^2^IA has been tested with major modern browsers such as Google Chrome, Safari, Mozilla Firefox and Microsoft Internet.

## Result

### Workflow

A standard data analysis workflow in M^2^IA contains six major steps: (1) data upload and processing, (2) global similarity analysis between microbiome and metabolome data matrix, (3) pairwise metabolite-microbe correlation analysis, (4) multivariate regression-based data integration analysis, (5) functional network analysis, and (6) an auto-generated html report for overview. Figure 1 summarizes the overall design and the flowchart of M^2^IA. Each step offers multiple methods helping users to explore complex correlations between microbiome and metabolome. In this section, each step will be described in detail.

**Figure 1.**
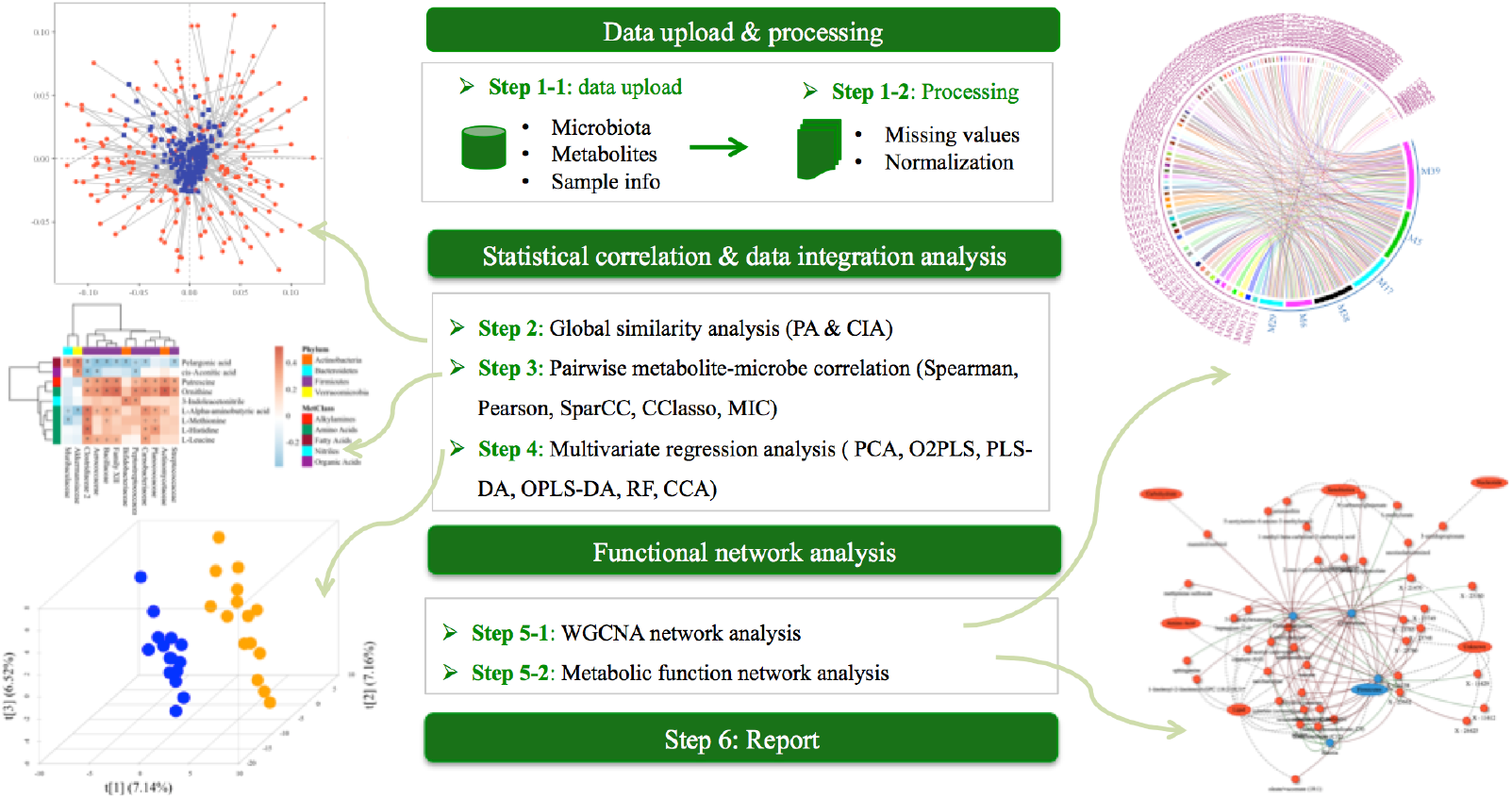
The flowchart of M^2^IA.

### Data upload and processing

#### Data upload

The first step is to upload three different types of data: (1) microbiome datasets: for 16s rRNA gene sequencing data, an OTU abundance table with taxonomic annotations (.txt format) and a corresponding reference sequence file (.fna format) are required. Similarly, a unigene table with taxonomic annotations and a table with KEGG KO function annotations are required for metagenomic shotgun sequencing study. (2) metabolome dataset: a metabolite abundance table (.csv format) with compound name, HMDB database ID, and chemical class information, and (3) a metadata table containing sample IDs and group information. Sample IDs from microbiome and metabolome datasets may not be exactly the same, but should be paired correctly within the metadata table. Once uploaded successfully, M^2^IA system examines the consistency of sample IDs from two datasets and notifies users whether there exist unmatched samples. After that, the metadata table is presented and can be modified by users interactively. For example, instead of uploading a new table, when users want to change grouping information or remove certain samples, they can modify group IDs or uncheck certain samples directly on the table for downstream data analysis.

#### Missing value processing

This step includes data filtering and missing value imputation. First, unqualified variables can be removed according to the criteria of sample prevalence and variance across samples. For microbiome data, the minimal count number, the percentage of samples with non-missing values, and the relative standard deviation (RSD) are used for screening. As suggested by MicrobiomeAnalyst^[10]^, features with very low count (e.g., <2) in a few samples (e.g., <20%), or very stable with limited variations across all the samples (e.g., RSD < 30%), are considered difficult to interpret their biological significance in the community and can be removed directly. Similarly, our previous work summarized that those metabolic features with missing values in more than 80% of samples or RSD values smaller than 30% can be filtered at the beginning. The sample prevalence is calculated based on all the samples or samples within each group^[13]^.

Second, both microbiome and metabolome have the characteristics of sparsity seen as the absence of many taxa or metabolites across samples due to biological and/or technical reasons^[14]^. Such missing values may pose general numerical challenges for traditional statistical analysis, thus M^2^IA provides a variety of methods for missing value imputation, aiming to remove zeros or NAs in the data matrix and facilitate subsequent statistical analysis. There are two different approaches to handle missing values: one is to simply replace missing values with a certain value (called pseudo count in microbiome), including half of the minimum or the median value; another is to apply regression models to impute missing values, such as random forest, k-nearest neighbors, Probabilistic PCA, Bayesian PCA, singular value decomposition, and the quantile regression imputation of left-censored data (QRILC)^[13]^. Users may choose an appropriate method for missing value imputation.

#### Data normalization

After missing value processing, users can perform normalization method in order to make more meaningful comparisons. M^2^IA provides the total sum scaling that calculates the relative percentage of features, or the log transformation when data does not follow normal distribution. To note, scaling or transformation methods are also provided for specific statistical analysis, such as principal component analysis (PCA) and PLS-DA analysis.

### Global similarity between two datasets

After data processing, the first step is to apply Coinertia analysis (CIA) and Procrustes analysis (PA) to evaluate the global similarity between metabolome and microbiome dataset (Figure 1)^[15]^. CIA can be considered as a PCA model of the joint covariances of two datasets. Similarly, PA measures the congruence of two-dimensional data distributions from superimposition and scaling of PCA models of two datasets (Figure 2A). The correlation coefficient R is between 0 and 1, and the closer it is to 1 indicates the greater similarity between two datasets.

**Figure 2.**
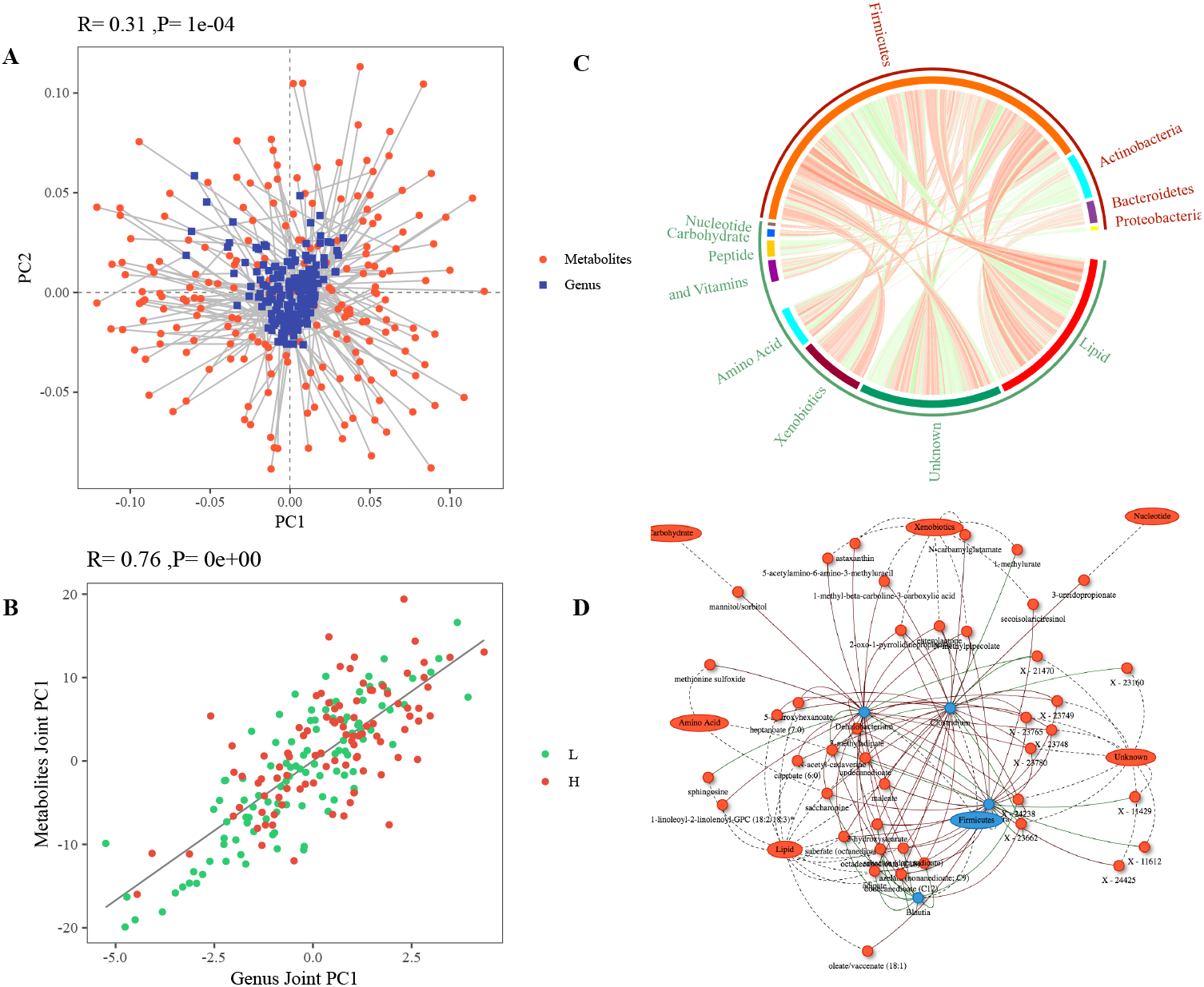
Illustration of similarity analysis and spearman correlation analysis results. (A) PA. The length of lines connecting two points indicates the agreement of samples between two datasets. (B) O2PLS model. (C) Circos plot of spearman correlations between microbes and metabolites. (D) Network of spearman correlations between differential metabolites and microbes belonging to the Firmicutes.

### Pairwise metabolite-microbe correlation analysis

#### Correlation analysis methods

M^2^IA provides five different types of correlation analysis, including Pearson, Spearman, SparCC, CCLasso, and Maximal Information Coefficient (MIC) analysis. Although each method has it pros and cons in different situations, users can choose an appropriate one. Since Spearman correlation analysis outperforms other methods due to their overall performances^[16]^, it has been set as a default one in M^2^IA. Microbes can be analyzed according to their different taxonomic annotations (i.e., phylum, genus, and species) and metabolites at different chemical classifications (e.g., amino acids, sugars, free fatty acids etc.). In addition, M^2^IA allows users to define the specific criteria of significance between microbe-metabolite pairs for subsequent analysis, e.g., absolute correlation coefficients > 0.3 and p value <0.05.

#### Visualization

Three different ways of visualization are provided in order to explore complex correlations between microbes and metabolites. The first one is to apply circos plot that can help users to quickly identify those microbes having close correlations with different classes of metabolites (e.g., amino acids) (Figure 2C). The second one is to apply a heat map to illustrate their relative positive/negative correlations between each microbe and metabolite. Meanwhile, hierarchical clustering is used to analyze the similarities among metabolites or microbes, in which closely correlated metabolites/microbes are usually clustered together. The third one is to provide a microbe-metabolite interaction network using cytoscape technique, which can help users to explore complex relationship between microbes and metabolites (Figure 2D).

### Multivariate regression–based data integration

#### Unsupervised multivariate analysis

M^2^IA provides canonical correlation analysis (CCA)^[17, 18]^ and O2PLS^[19]^ in order to evaluate the inherent correlations between two datasets without considering phenotype information. CCA aims to find two new bases (canonical variate) in which the correlation between original parameters of two datasets is maximized. O2PLS is capable of modeling both prediction and systematic variation, and the joint score plot indicates their relationship between two data matrix. Metabolites/microbes with the large canonical coefficients from CCA model or loading values from O2PLS model are considered essential ones for their similarities (Figure 2B).

#### Supervised multivariate analysis

In comparison, supervised multivariate analysis methods integrate two data matrix initially and identify differential variables (microbes & metabolites) that significantly contribute to the discrimination of different groups. Here, M^2^IA offers PCA score-based differential analysis, partial least squares discriminant analysis (PLS-DA), orthogonal partial least squares discriminant analysis (OPLS-DA), and random forest (RF). PCA is originally an unsupervised data mining method; here we examine the differences of the first principal component score values from PCA model, and their correlations with the phenotype information. PLS-DA and OPLS-DA have been commonly applied in the field of metabolomics for data dimension reduction and feature selection. OPLS-DA seeks to maximize the explained variance between groups in a single dimension or the first latent variable (LV), and separate the within group variance (orthogonal to group difference) into orthogonal LVs (Figure 3A). The variable loadings and/or coefficient weights from a validated PLS-DA and OPLS-DA model are used to rank variables with respect to their performance for discriminating between groups (Figure 3B). Boruta algorithm-based RF classifier can identify important features by shuffling samples and adding extra randomness to the system (Figure 3C).

**Figure 3.**
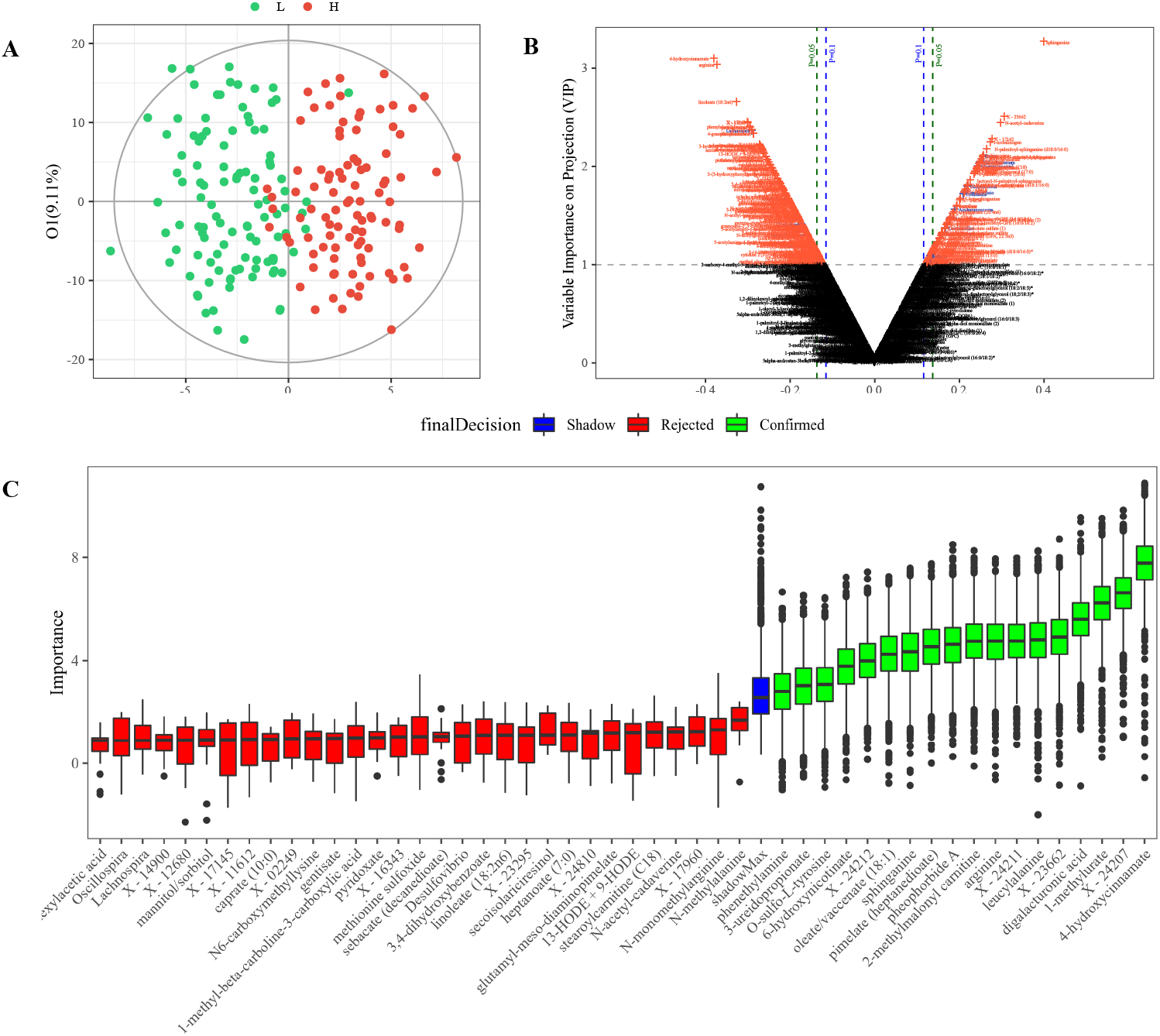
(A) Score plot of an OPLS-DA model between OB and HC group. (B) Variable importance plot (V-plot) of an OPLS-DA model between OB and HC group. (C) Feature selection of Random Forest modeling.

### Network analysis

#### WGCNA-based network analysis

Weighted gene co-expression network analysis (WGCNA) was recently raised to integrate high-throughput ‘-omics’ datasets for identifying the potential mechanistic links^[11]^. In M^2^IA, WGCNA algorithm is used to collapse co-abundant metabolites into different clusters, and thus metabolites within a cluster are highly correlated. One attractive feature is that both identified and unknown metabolite features can be considered. KEGG microbiome functional modules are annotated using Tax4fun2^[20, 21]^ for 16s RNA sequencing data or HUMAnN pipeline^[22]^for shotgun sequencing data, respectively. Finally, the correlations between metabolite clusters and microbial functional modules with phenotype information are examined using Spearman correlation analysis and further visualized using heat map and network.

#### Metabolic function analysis

The final step aims to analyze or predict the biological function profiles using metabolome and microbiome data, respectively. The metabolic pathway activity is analyzed using PAPi package in R software ^[23]^. In parallel, function profiles of microbial communities are predicted using Tax4Fun tool for 16s rRNA data and HUMAnN for shotgun sequencing data. Linear discriminant analysis is further used to identify differential KO identifiers from microbiome and metabolome (Figure 4A-C). Their relationships are further investigated using a microbe-metabolite interaction network by connecting a specific metabolite, its reactions and related enzymes, and the genomes of organism (both host and microbial)(Figure 4D-E).

**Figure 4.**
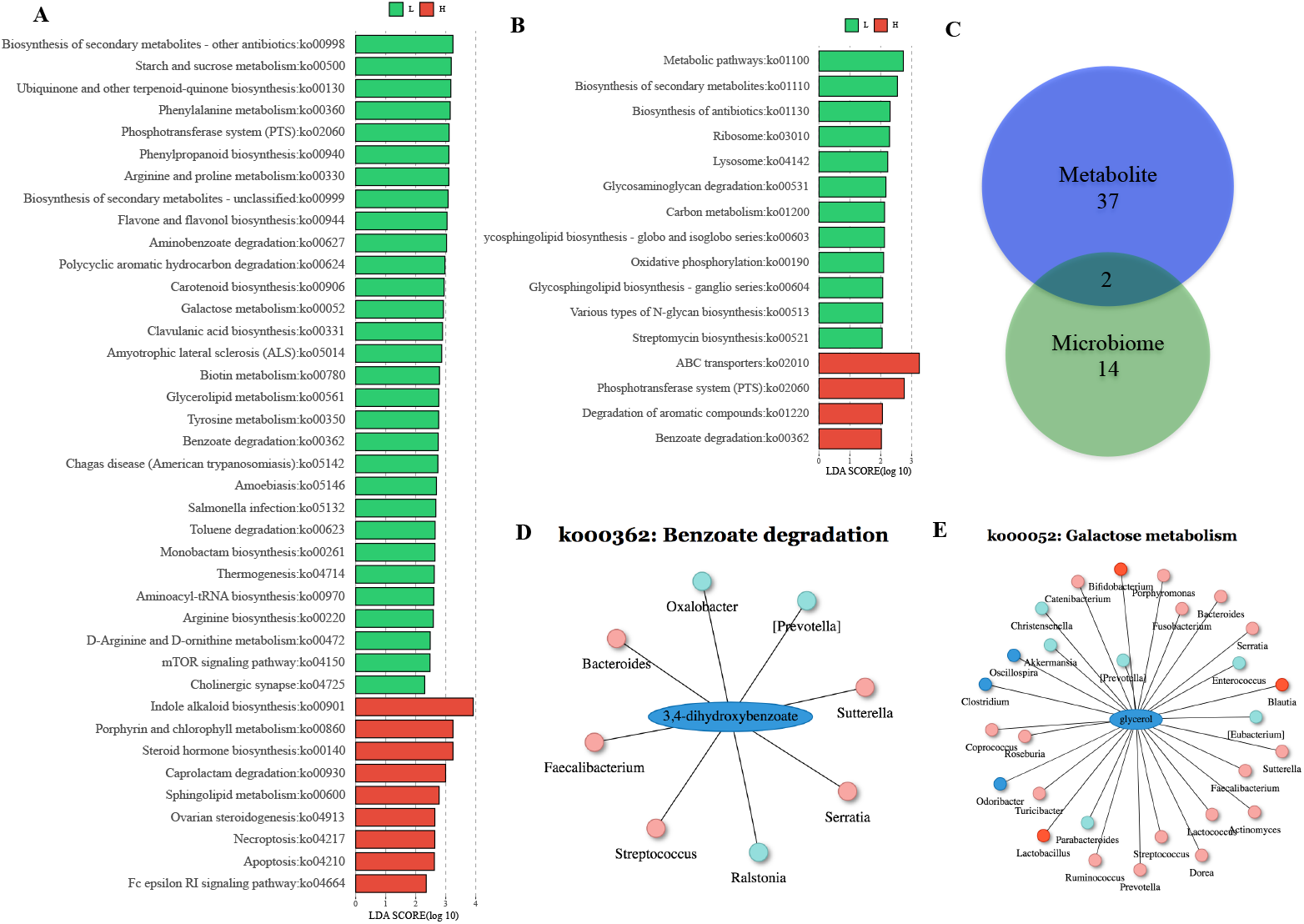
(A-B) Bar plot of differential KO functions from metabolome and microbiome data. (C) Venn diagram of differential KOs. (D-E) Interaction network of each metabolite (3,4-dihydroxybenzoate and glycerol) with microbes that may participate its metabolism.

### Automated html report

Once users upload the data and optimize parameters of data analysis, M^2^IA provides a simplified user experience through a one-click button to submit a job. It may take a few minutes for M^2^IA to perform data analysis and generate interpretable results. The exact processing time depends on the sample size, number of variables, and number of pairwise group comparisons. Once the analysis is completed, M^2^IA will summarize key results and produce an html report for users. In addition, a zip-compressed package containing all the supporting data files (tables and figures) can be downloaded for future use.

### Application example

To validate the performance of M^2^IA, we analyzed the fecal metabolome and microbiome of 786 individuals from TwinsUK cohort^[24]^. 16S rRNA was extracted from fecal samples, PCR amplified, barcoded per sample and sequenced with the Illumina MiSeq platform, as previously described^[25]^. A total of 1,116 different metabolites were measured using UPLC-MS/MS platforms. First, the Procrustes Analysis revealed moderate similarities between metabolome and microbiome with statistical significance (Figure 2A, r=0.31, p<0.05). Since most of individuals at the metabolic level surrounded those at the microbiome level, the variations of metabolic profiles among individuals were larger in metabolome. The O2PLS analysis indicated microbial and metabolic data matrixes were highly correlated with correlation coefficient 0.76 (p<0.05, Figure 2B), and those variables that contributed to their similarities could be identified according to their weights. Then, Spearman correlation analysis was performed to explore specific microbe-metabolite correlations. The circos plot showed seven different classes of metabolites (i.e., nucleotides, carbohydrates, peptides, cofactors and vitamins, amino acids, xenobiotics, and lipids) were correlated with microbes mainly belonging to Firmicutes, Actinobacteria, Bacteroidetes, and Protebacteria (Figure 2C). Among which, four microbes at genus level belonging to the Firmicutes had the most significant correlations with metabolites, and their specific correlations were further visualized in a network (Figure 2D).

According to BMI, participants were divided into obese (OB, BMI >28) and healthy control group (HC, BMI <24). The multivariate OPLS-DA model based on the integrated data matrix of metabolome and microbiome indicated the obvious difference between OB and HC group (Figure 3A). Differential metabolites and microbes contributing to their separations were identified using both VIP values (VIP>1) and correlation coefficients (p<0.05), as indicated by the V-plot (Figure 3B). Alternative multivariate models e.g., RF model, could be further used to validate the relative contributions of microbes and metabolites to the discrimination between two groups (Figure 3C). For example, arginine was identified as a significant marker from both OPLS-DA and RF model. Then, we performed functional network analysis to further interpret the biological significance of their correlations. A total of 39 and 16 significant biological functions (KOs) were identified in microbiome and metabolome between OB and HC group, respectively (p < 0.05, Figure 4A,B). Two important pathways (i.e., the benzoate degradation and phosphotransferase system) were overlapped (Figure 4C), among which, the benzoate degradation was the most significant metabolic pathway in both microbial and metabolic function analysis. Then, each differential metabolite and relevant microbes that may participate in its metabolic reactions were further identified through searching the KEGG database. For example, 3,4-dihydroxybezoate (also called protocatechuic acid) in the benzoate degradation was a major metabolite of antioxidant polyphenols formed by gut microbiota, and might have beneficial effects against inflammation and insulin resistance in obesity^[26]^. Our results indicated that 3,4-dihydroxybezoate was significantly reduced in obesity, and eight different gut microbes might participate in its metabolism (Figure 4D). Similarly, a total of 28 different microbes identified in this study were involved in the glycerol metabolism of phosphotransferase system (Figure 4E).

## Discussion

We have implemented M^2^IA pipeline as a user-friendly web server that can provide microbiome and metabolome data integration analysis to understand the important roles of microbial metabolism in diverse disease contexts. This is a bioinformatics workflow that integrates a wide range of univariate and multivariate methods, including PA/CIA-based data matrix similarity analysis, univariate-based correlation analysis, multivariate regression-based analysis, and knowledge-based function analysis. The advantage of applying these methods simultaneously is to understand the inherent characteristics of the high-dimensional omics data in different ways, and obtain reliable results from multiple analyses. M^2^IA also implements KEGG database in order to link orthologous gene groups to metabolic reactions and annotated compounds. So that researchers will have better understanding of their origins of metabolites, from host or microbial, and what microbes participate in specific metabolic reactions. To make it more convenient, M^2^IA automatically produces a comprehensive report that summarizes and interprets the key results.

Users can apply M^2^IA to their own microbiome and metabolome data. In terms of analytical platforms, M^2^IA accepts the microbiome data from 16S rRNA gene sequencing or shotgun metagenomic sequencing technologies, and metabolomics data from mass pectrometry or NMR analytical platforms. For sample types, many human studies on gut microbiota collect fecal samples, due to its non-invasive characteristics. The fecal metabolome provides a functional readout of the gut microbiome, and the integrative analysis can provide better understanding of correlations between gut microbiome and metabolome. We recommend applying the same fecal samples of human subjects or fecal/colon contents of animals for analysis. However, samples types can be from any site, i.e., saliva, buccal mucosa, and colon tissue samples. Simultaneously, metabolomics studies can analyze feces, plasma/serum, urine, saliva, exhaled breaths, cerebrospinal fluid, and tissues of target organs. Thus, sample types may differ, but they need to require originating from a same subject for both microbiome and metabolome analysis.

Finally, the limitations and future directions of this study disserve to be mentioned. M^2^IA accepts annotated microbioa and metabolites as input data matrix. However, the raw data preprocessing steps are not included in this pipeline, e.g., quality control and microbial annotation for microbiome data, and peak detection and metabolite annotation for metabolomics data. Currently, many sophisticated software tools or pipelines have been developed for original data preprocessing, such as Qimme2^[27, 28]^ for 16s RNA sequencing, and XCMS for MS-based metabolic profiling^[29]^. M^2^IA focuses on providing a standardized pipeline and a comprehensive workflow for the integrative analysis of metabolome and microbiome data, not only exploring the statistically significant correlations but also investigating the biological significance of their interaction network. Moreover, KEGG database based functional network analysis can help to explain their biological correlations and develop mechanistic hypothesis potentially applicable to the development of nutritional and personalized therapies. In the future, more advanced integration methods will be introduced in M^2^IA and more independent knowledge databases (e.g., enzyme database and drug metabolism database) will be integrated within M^2^IA for deeper biological interpretation.

## Conclusions

In summary, M^2^IA is an effective and efficient computational tool for experimental biologists to comprehensively analyze and interpret the important interactions between microbiome and metabolome in the big data era.

## Authors’ contributions

YN contributed to develop the methodology, algorithms, and write the manuscript. YG contributed to the software development and implementation. YD provided valuable testing datasets during method development of our pipeline. PW and CS contributed to TwinsUK data preparation and processing for pipeline validation. HC and FJ provided valuable suggestions on software development and manuscript modification. FF supported this project and contributed to the review and modification of the manuscript.

## Availability of data and materials

Data used in this study can be accessed at http://m2ia.met-bioinformatics.cn.

## Acknowledgements

Not applicable

## Ethics approval and consent to participate

Not applicable

## Consent for publication

Not applicable

