## Supporting information for "M^2^IA: a Web Server for Microbiome and Metabolome Integrative Analysis"

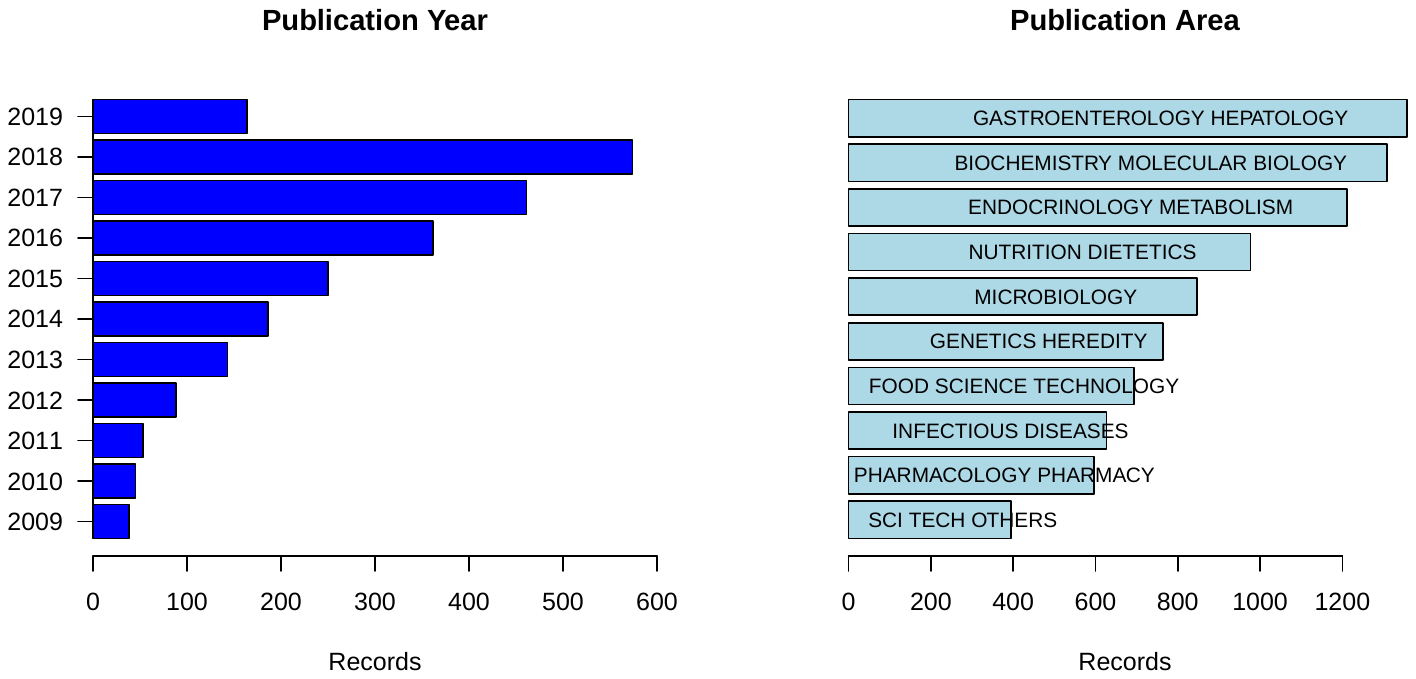

**Figure S1**. Analysis of 2,630 publication records using TITLE: (microbio*) AND TITLE: (metabol*) in the Web of Science database searched on March 23 2019. (A) The number of records from 2009 to 2019. (B) The list of top ten publications area.

Table S1. Example studies for integrative analysis of microbiome and metabolome

| Authors | Diseases | Species | Number | Microbome Sample type | Microbiome Platform | Metabolome Sample type | Metabolome Platform | Integrative methods |
| --- | --- | --- | --- | --- | --- | --- | --- | --- |
| Ian H McHardy et al | Health | Human | 47 | fecal | 16S | fecal | UPLC | Spearman |
| Barton et al | Health | Human | 86 | fecal | 16S | urine, fecal | H-NMR,RP,HILIC,UPLC-MS,GC-MS | Spearman, OPLS-DA, PCoA, PCA |
| Tianlu Chen et al | Health | Rat | 294(42) | fecal | 16S | brain, blood, fecal | UPLC/QTOF-MS and UPLC/TQ-MS | Spearman, rCCA,network |
| Zierer J et al | Health | Human | 786 | fecal | 16S | fecal | LC/MS | GeneNet package |
| Dhakan DB et al | Health | Human | 110 | fecal | 16S | serum, fecal | GC-MS | Spearman |
| Jing Li et al | Hypertension | Human | 196 | fecal | 16S | fecal | LC/MS | Spearman, PLS-DA, OPLS-DA |
| Xiao Cui et al | chronic heart failure | Human | 94 | fecal | Metagenomome | fecal, plasma | LC/MS | Spearman |
| Aleksandar D. Kostic et al | T1D | Human | 989(33) | fecal | 16S | fecal, serum |  | Spearman (penalized canonical correlation analysis) |
| Tiejuan Shao et al | Gout | Human | 52 | fecal | 16S | fecal | 1H-NMR | Spearman |
| Christopher J Stewart et al | Bronchiolitis | Human | 144 | nasopharyngeal mucus | 16S | nasopharyngeal mucus | UPLC-MS | Spearman |
| Ruixin Liu et al | Health(obesity) | Human | 257 | fecal | Metagenomome & 16S | serum | HPLC-MS,UHPLC | Spearman, CIA, CCA |
| Mohammad Sajjad Ghaemi et al | Health(pregnant) | Human | 68 | vaginal swabs, stool, saliva gum | 16S | plasma | LC-MS | Spearman |
| Antoine M. Snijders et al | Health | Mouse | 30 | fecal | 16S | fecal | GC-MS | MIMOSA |
| Tanja V Maier et al | Health | Human | 39 | fecal | 16S | fecal | FT-ICR-MS | Context likelihood of relatedness (CLR) |
| Elena Sanguinetti et al | Health | Mouse | 33 | fecal | 16S | fecal, serum | 1H-NMR | Spearman, Pearson |
| Federica Del Chierico et al | NFLD | Human | 115 | fecal | 16S | fecal | VOCs | Spearman, PCA ,PLS, PLS-DA |
| Maria Laura Santoru et al | IBD | Human | 132 | fecal | 16S | fecal | GC-MS,H-NMR,LC-QTOF-MS | Spearman |
| Yang Liu er al | liver cirrhosis | Human | 70(28) | fecal | 16S | fecal | UPLC/MS | Spearman |
| Cyrielle Caussy et al | NAFLD | Human | 156 | fecal | Metagenomome | serum | GC/MS,LC/MS/MS | Spearman |
| Eric A Franzosa et al | IBD | Human | 220 | fecal | Metagenomome | fecal | LC–MS | Spearman |
| Stewart, Christopher J et al | Sepsis | Human | 613(35) | fecal | 16S | fecal | UPLS-MS | (sparse partial least squares regression) sLPS |
| J. S. Bajaj et al | minimal hepatic encephalopathy | Human | 30 | fecal | Pyrosequencing | fecal, serum, urine | HPLC,GC/MS,LC/MS |  |
| Casey M Theriot et al | Health | Mouse | 80 | fecal | 16S | fecal | GC-MS | Spearman |
